# Deep data mining reveals variable abundance and distribution of microbial reproductive manipulators within and among diverse host species

**DOI:** 10.1101/679837

**Authors:** Paloma Medina, Shelbi L Russell, Kavya Aswadhati, Russell Corbett-Detig

## Abstract

Bacterial symbionts that manipulate the reproduction of their hosts are important factors in invertebrate ecology and evolution. Studying the genomic and phenotypic diversity of reproductive manipulators can improve efforts to control infectious diseases and contribute to our understanding of host-symbiont evolution. Despite the vast genomic and phenotypic diversity of reproductive manipulators, only a handful of strains are used as biological control agents because little is known about the broad scale infection frequencies and densities of these bacteria in nature. Here we develop a data mining approach to quantify the number of arthropod and nematode host species infected with *Wolbachia* and other reproductive manipulators such as *Rickettsia* and *Spiroplasma*. Across the entire Sequence Read Archive (SRA) database, we found reproductive manipulators infected 2,083 arthropod and 119 nematode samples, representing 240 and 8 species, respectively. After accounting for sampling and infection frequency differences among species, we estimated that *Wolbachia* infects approximately 44% of all arthropod species and 34% of all nematode species. In contrast, we estimated other reproductive manipulators infect 1-8% of arthropod and nematode species. Next, we explored another important biological parameter: the relative bacterial density, or titer, within hosts. We found variation in titer within and between arthropod species to be large, and that host species explains approximately 36% of variation in titer across our dataset. This suggests bacterial strain and/or host species plays a role in shaping bacterial densities within and between host species. By leveraging the model system *Drosophila melanogaster*, we also found a number of host SNPs associated with titer in genes potentially relevant to host interactions with *Wolbachia*, suggesting bacterial induced host genome evolution. Our study demonstrates that data mining is a powerful tool to understand host-symbiont co-evolution and opens an array of previously inaccessible questions for further analysis.

## Introduction

Bacterial symbionts of eukaryotic hosts exhibit an impressive array of phenotypes that interact with host biology. Included among these symbionts are bacteria that alter host reproduction in order to increase their likelihood of transmission to the next host generation [1–4], a strategy termed reproductive manipulation. Depending on the nature of host reproduction, reproductive manipulators are generally transmitted vertically by associating with either the oocyte or developing embryos [5]. Manipulative phenotypes range from strategies that prevent the survival of uninfected offspring, such as cytoplasmic incompatibility, to strategies such as feminization, male killing, and parthenogenesis, that actually change the sex ratio to favor females for overall increased infection rates [1–4]. These drastic manipulations of normal host biology mean that reproductive manipulators can induce reproductive isolation, drive changes in sexuality, and alter the reproductive ecology of their hosts [6–8].

While reproductive manipulators undergo vertical transmission to reach the next host generation, many species and strains also exhibit low rates of horizontal transmission between contemporary host species [5]. This combination of transmission strategies coupled with reproductive manipulation serves to quickly increase the prevalence of reproductive manipulators in a host population and to spread them rapidly among host species.

Given the impressive abilities of these bacteria to colonize new hosts and impact host fertility and development for their own reproduction, much effort has been spent attempting to estimate the infection frequencies of reproductive manipulators. Due to its wide distribution and presence in model organisms such as *Drosophila, Wolbachia* is one of the most well studied symbionts in general, and is the best studied reproductive manipulator in particular. Estimates of the frequency of *Wolbachia* infections amongst arthropods range from 11% [9] to 76% [10]. The large variation in estimates is likely an effect of methodological differences including sampling bias and variation in PCR assay sensitivity. Other reproductive manipulators in the *Cardinium, Arsenophonus, Rickettsia*, and *Spiroplasma* clades are reported to occur in 4% to 7% of all species [11]. However, these estimates are even less certain than those for *Wolbachia* because less infection frequency data is available for these clades, and similar sampling and assay biases might impact frequency estimates. Given that the probability of sampling an infected individual is positively correlated to the prevalence of the symbiont and the number of individuals sampled from the host population, undersampling also imposes a barrier to confident detection of low frequency infections.

Reproductive manipulators also experience a range of fitness tradeoffs during host growth and development. A symbiont must be present at sufficiently high frequencies within a host to promote successful transmission to subsequent host generations. However, exceedingly high abundance of symbiont cells relative to host cells, termed titer, may impose a significant fitness cost for the host and their symbionts [12, 13]. Reproductive manipulators may be more virulent to their hosts when they achieve high titers, and natural selection on symbiont proliferation and host regulation should favor the evolution of intermediate symbiont frequencies [14, 15]. This tradeoff between reliable transmission and fitness costs to hosts is therefore an essential component of understanding reproductive manipulator co-evolution. Nonetheless, little is known about the relative abundances of reproductive manipulators within host individuals in large part due to the challenges of collecting these data in high throughput ways.

Whole genome sequencing and bioinformatic approaches offer appealing alternatives to conventional PCR-based survey methods to estimate reproductive manipulator infection rates. These methods are not biased by primer selection, are less sensitive to false positives due to contamination, and enable testing of large numbers of samples. Similarly, genome-sequencing is a potentially powerful tool for interrogating symbiont titer within host individuals. With Illumina shotgun sequencing, when the genome of a potential host individual is sequenced, the host’s symbionts are also sequenced. This makes the publicly-available databases a treasure trove for sampling reproductive manipulators with bioinformatic approaches. In fact, the NCBI Sequence Read Archive (SRA) [16] contains ∼60,000 sequencing runs for samples tagged as Arthropoda alone. Searching, or mining, genomic sequencing data has been shown to be a cost effective and powerful strategy to detect *Wolbachia* infections [9, 17]. However, prior studies did not include other reproductive manipulators [9, 17] or were focused on a single host species [17]. Indeed, one of the most compelling arguments for using non-targeted, publicly available data is to mitigate ascertainment biases towards selecting species already known to harbor reproductive manipulator infections.

Characterizing the prevalence and distribution of reproductive manipulators could be especially valuable to biomedical researchers using reproductive manipulators to control the spread of human pathogenic viruses such as Zika, Dengue and Chikungunya [18–20]. Currently, only the wMel strain of *Wolbachia* is being used as a biological control agent [20, 21]. However, the large genetic and phenotypic diversity of reproductive manipulators could suggest different species or strains of bacteria to more effectively combat the spread of arboviruses. Additionally, a strategy that opportunizes on cytoplasmic incompatibility, termed the “sterile male technique”, can be used to control mosquito population sizes [22, 23]. As this strategy requires low infection frequencies in females to work [24], and different strains may have compatible rescue abilities [25], it is necessary to understand the diversity and distribution of these bacteria in nature. Cataloguing the distribution and titer of reproductive manipulators builds a foundation to explore reproductive manipulators as powerful and versatile biological control agents.

Here, we establish and validate a computational pipeline to determine the infection status of host samples downloaded from the SRA database and estimate the titer of symbionts within positively infected host individuals. We use this approach to (i) quantify *Wolbachia* infection frequency and infection titer of hosts, (ii) quantify the infection frequency and titer of other genera including a range of reproductive manipulator symbiont clades, (iii) compare infection frequencies and titers (from i and ii) among host groups, and (iv) identify *Drosophila melanogaster* SNPs associated with *Wolbachia* titer. From this work, we classified 2,083 arthropod and 119 nematode samples as infected with a reproductive manipulator, including 95 species with previously unknown infections. Additionally, we show substantial variation in symbiont titer within and between arthropod host species. Moreover, we show that symbiont titer does not vary systematically across bacterial clades which indicates that other evolutionary, ecological, or physiological processes may explain the disparate global distributions of reproductive manipulators. We present extensive validation of our methods, including orthogonal *in vivo* approaches, to help validate our large-scale data mining approach for addressing fundamental questions in symbiont genomics and evolutionary biology.

## Methods

### Reproductive manipulator reference genome panel

We built our BLAST database using RefSeq genome assemblies for *Arsenophonus, Spiroplasma, Rickettsia*, and *Wolbachia*. We included three *Arsenophonus*, six *Wolbachia*, 27 *Spiroplasma*, five *Cardinium*, and 61 *Rickettsia* genome assemblies (Supplementary Table S1). These genomes were selected to span the known diversity of these bacterial groups. On average, each genome assembly was about 1.3 Mb. We used these 102 genome assemblies to build a BLAST database using the blastdb command from the NCBI blast package (version 2.7.1). We also included additional *Wolbachia* reference genomes to estimate symbiont titer, which are listed in Supplementary Table S1, bringing the total number of references to 141.

### Arthropod and nematode SRA dataset

We downloaded all Arthropoda sequencing read data from the NCBI Sequence Read Archive (SRA) database [16]. We filtered samples under the Arthropoda and Nematoda taxonomy for those sequenced on Illumina sequencing platforms. We filtered nominally for whole genome shotgun libraries, but for completeness we further removed samples that were marked as “reduced representation”, “chipseq”, and other terms that preclude a fully random, shotgun library approach. In total, we tested 27,256 arthropod and 5,229 nematode samples for reproductive manipulator infections. We consolidated all subspecies into a single species which resulted in a total of 1,299 arthropod and 128 nematode species. SRA metadata for samples we classified can be found in Supplementary Tables S2 and S3.

### Determining Infection Status using a BLAST-based approach

In order to classify a host sample as positively or negatively infected with a reproductive manipulator, we analyzed local alignments between host DNA sequence data and reproductive manipulator reference genomes. We binned each reference genome into 5kb segments and computed the proportion of bins with a significant hit (breadth of coverage). We also computed the variance of BLAST hits across each bin and estimated the coverage of the reproductive manipulator genome (Supplementary Method S1). We determined a sample to be a candidate for a positive infection if it had a 90% breadth of coverage and >1x estimated coverage on a reproductive manipulator reference genome.

### Validation of bacterial detection pipeline

To estimate the sensitivity and the specificity of our reproductive manipulator annotation method, we compared our results to a previous extensive survey conducted by [17], which was PCR validated. We determined the reproductive manipulator infection status of 158 individuals from the *Drosophila Genetics Research Panel* (DGRP) [26] using our computational pipeline. These 158 *D. melanogaster* samples had matching PCR and WGS determined infection statuses generated by [17] and had their sequence data available on the NCBI SRA database. In addition, there were 16 samples we excluded from our analysis because four of them were not available on the SRA, eight did not have corresponding PCR and WGS *Wolbachia* infection statuses, and four had discrepancies in infection status between sequence iterations of DGRP lines (Supplementary Method S2). See Supplementary Method S2, Table S4, and Table S5 for subsampling validation methods, pipeline accuracy to divergent reference strains, and a comparison of our approach to other methods [9].

### Beta-binomial estimation of reproductive manipulator prevalence

Beta-binomial distributions have been used to fit *Wolbachia* infectious status across species previously [27]. The beta-binomial model considers N random variables, X_j_, which are all binomially distributed (*i*.*e*., infected vs. not-infected), but each with different parameters q_j_ and n_j_, so that X_j_∼Bin(q_j_, n_j_) (Figure S1). Using the approach developed in [27], we determined (1) moment estimators *u* and *s*, (2) beta distribution parameters α and □, and (3) the global infection rate *x*. After we fit a beta distribution to the data, we took the integral from *c* to 1, where *c* is the minimum infection rate of a species to be considered positively infected. For example, a value of 0.001 means if one individual in 1000 is classified as positive, the species would be classified as being positively infected. For consistency with previous work, we set *c* to equal 0.001 (as in [27]). See Supplementary Method S3, Figure S2, and Table S6 for beta-binomial rationale and downsampling results, which aimed to reduce sampling intensity bias.

## Estimating Symbiont Titer

We estimated the ratio of symbiont genome compliments to host genome compliments, hereafter referred to as titer (Supplementary Method S4). First, we computed the number of symbiont genome compliments through our BLAST-based method with all available *Wolbachia, Arsenophonus, Spiroplasma*, and *Rickettsia* genomes available on NCBI (Supplementary Method S1, Supplementary Table S1). Next, we computed the number of host genome compliments by aligning DNA sequencing reads to single copy orthologous arthropod proteins from OrthoDBv9 [28] and taking the average of maximum depth across proteins with hits. We report the symbiont haploid : host pseudo haploid titer computation throughout this manuscript. See Supplementary Methods S4 and Figures S3 and S4 for validation of our approach to estimate host genome compliments.

### Drosophila oocyte sampling, imaging, and analysis

We obtained *Drosophila melanogaster* and *Drosophila simulans* fly stocks infected with the wMel and wRi strains of *Wolbachia* from the Sullivan Lab. Stage 9/10a oocytes were dissected from these flies, stained and mounted on glass slides, and imaged with a SP5 Leica confocal microscope. We analyzed the fluorescence due to *Wolbachia* as described in [29] (See Supplementary Method S5).

### Estimating the Biological and Methodological Contributions to Titer Variation

We fit several models to determine how much host/symbiont genetics contributes to variation in titer. We fit Generalized Linear Models models to log transformed arthropod titer data using the glm() functions in R version 3.5.0. Specifically, we fit a GLM of the form: glm(log(titer) ∼ arthropod_species). We also compared that model to one including reproductive manipulator clade (e.g. “*Wolbachia*”) using a GLM of the form: glm(log(titer) ∼ arthropod_species + reproductive_manipulator_clade. We used a likelihood ratio test to compare these two model fits using anova(model1, model2, test=“Chisq”). In addition, to test for an effect of sequencing strategy (see also Supplementary Method S8), we fit a GLM of the form: glm(log(titer) ∼ arthropod_species + pool_status).

## Software Availability

The pipeline and individual scripts used to classify microbial reproductive manipulator infections and estimate titer can be accessed through GitHub (www.github.com/pamedina/prevalence).

## Results and Discussion

### Reproductive Manipulators Infect Arthropod and Nematode Hosts in the SRA

We developed a powerful bioinformatic pipeline to identify reproductive manipulator infections within sequencing datasets. Briefly, our approach compares sequencing reads from a given sample to a set of reference genomes from reproductive manipulator species to determine if a given host is positively infected (Figure 1, see Methods). We extensively characterized the sensitivity and accuracy of our bioinformatic pipeline using previously known *Wolbachia* infection statuses of the Drosophila Genetic Reference Panel (DGRP [26, 30]) and other samples known to harbor genetically divergent *Wolbachia* [31] (see Supplementary Method S2). Using the confident infection status lines in the DGRP as our validation set, our method is completely concordant with both PCR and previous bioinformatic methods. Additionally, our approach remains accurate even for references that exhibit 5-15% pairwise sequence divergence (Supplementary Tables S7 and S8, Figures S5 and S6), indicating that our pipeline is an accurate and robust method to determine reproductive manipulator infections in the majority of host samples.

**Figure 1.**
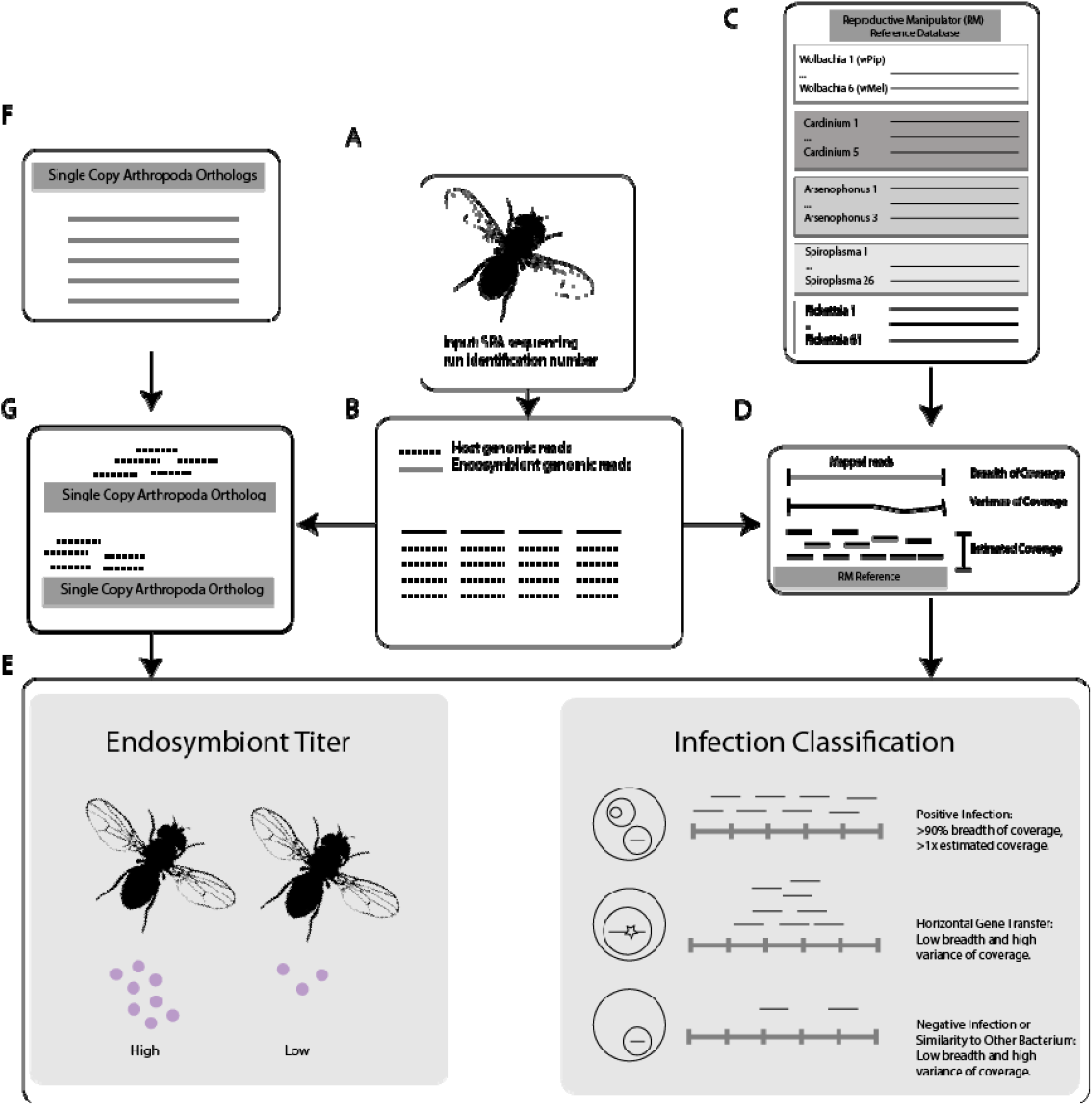
A schematic of the computational pipeline used to determine the reproductive manipulator infection status and endosymbiont titer of a sequencing run (see also Supplementary Methods S1 and S7). The pipeline **(A)** takes in a sample’s unique identification number, then **(B)** downloads two million endogenous and exogenous reads. Then, **(C)** reads are aligned to *Wolbachia, Arsenophonus, Spiroplasma, Cardinium*, and *Rickettsia* reference genomes, and **(D)** summary statistics for the sample aligned to each reference are computed. If a sample had between 0.1 and 0.9 breadth of coverage, the full dataset was downloaded and the workflow repeated to prevent false negative calls. **(F and G)** To estimate host coverage without requiring a reference genome, reads are also aligned to a set of 1066 single copy ancestral orthologs obtained from ORTHODB v9. Then, **(E)** we apply coverage breadth and depth cutoffs to classify infection status as positive, or negative. We compare the coverage of host reads to single copy orthologs to the coverage of endosymbiont reads to reproductive manipulator reference genome to estimate endosymbiont load.

Using our bioinformatic classification pipeline (see Methods, Figure 1), we tested nearly all arthropod and nematode samples on the SRA database that had been whole genome shotgun sequenced (as of January 22, 2020) and found 240 arthropod species and 8 nematode species had samples infected with a reproductive manipulator out of 1,299 and 128 species tested, respectively (Table 1, Supplementary Figure S7 and S8, Table S9 and S10, File S1 and S2). We identified 95 arthropod species with previously unreported reproductive manipulator infections (Supplementary Table S11 and Method S6).

**Table 1.** Raw infection counts of *Wolbachia, Spiroplasma, Rickettsia* and *Arsenophonus* infecting arthropods and nematodes. The frequency of positive samples or species is listed in parentheses.

|  | Samples |  | Species |  |
| --- | --- | --- | --- | --- |
|  | Arthropoda | Nematoda | Arthropoda | Nematoda |
| **Total** | 27256 | 5229 | 1299 | 128 |
| **Infected With Any RM** | 2083 (0.08) | 119 (0.02) | 240 (0.18) | 8 (0.06) |
| **Infected With**<br>***Wolbachia*** | 1978 (0.07) | 119 (0.02) | 179 (0.14) | 8 (0.06) |

*Wolbachia* was the most frequent reproductive manipulator in arthropod samples and species (1,978 samples, 179 species, Table 1). We also found numerous arthropod species infected with other clades of reproductive manipulators (Supplementary Table S9). Specifically, *Cardinium, Rickettsia, Spiroplasma*, and *Arsenophonus* infect two, 54, 69, and nine arthropod species in our dataset, respectively. The substantially lower infection rates for other reproductive manipulators than for *Wolbachia* are consistent with prior work [32–34] (Supplementary Table S10, Figure S8).

*Wolbachia* was the sole reproductive manipulator found in nematodes besides an instance of *Cardinium*, as expected based on previous work [11, 35]. Almost all nematode species infected with *Wolbachia* are filarial worms (Supplementary Table S10), a result supported by previous studies showing that filarial nematodes and *Wolbachia* are in an obligatory, mutualistic relationship [36–38]. There was one non-filarial nematode species infected with *Wolbachia*: *Pratylenchus penetrans*, a plant-parasitic nematode (order Tylenchida). This species has been shown previously to be infected with *Wolbachia* [35, 39]. These results, in addition to our validation experiments, indicate that our approach can accurately detect symbiont infections.

We analyzed co-infections at two different levels: species harboring samples infected with more than one reproductive manipulator clade and single samples infected by more than one bacterial clade. We found high rates of co-infections of different reproductive manipulators among arthropod hosts species. Species co-infections occurred in 29 of the 240 arthropod species that harbored any infection (*p* < 0.001 permutation test, Supplementary Method S7, Figure S9, and Table S12). Our observed rate of co-infection is consistent with previous work that identified eight out of 44 species were co-infected ([11], *p* = 0.34, Fisher’s Exact Test). However, we found no evidence for an excess of individual samples infected by more than one reproductive manipulator relative to expectations obtained by permutation (*p* = 0.155, 27 individuals, Supplementary Figure S10). These results may indicate that a subset of hosts, at the species level, are more likely to acquire symbionts than others or that they are more permissive to prolonged infections. This is supported by the observation that some host species harbor genetic variation that influences the abundance of symbionts within individuals [14, 40]. Alternatively, any direct or indirect mutualism between symbionts might facilitate the build-up of reproductive manipulators within a single species or host maternal lineage.

### Global Rates of Infection Range from 1-44% Across Symbiont Taxa

Our study aims to estimate the infection rate of reproductive manipulators in arthropod and nematode species overall, however, different taxa have been sampled to varying degrees and infection frequencies vary (Supplementary Table S9) so the number cannot be directly calculated from infection counts of individuals. To evaluate and correct for this ascertainment bias, we use an approach based on modelling observed infections using a beta-binomial distribution to estimate the total proportion of species infected with a reproductive manipulator bacterial species as has been done previously [27] (Supplementary Method S3, Figure S1). Using this approach, we estimated that 44% (95% CI 29-61%), and 34% (95% CI 5-69%), of arthropod and nematode species, respectively, are infected with *Wolbachia* (Table 2). We note that these values are consistent with expectations from previous work [10, 41–47]. Moreover, because the SRA has been populated with samples mostly without considering *Wolbachia* infection status, our approach should provide a relatively unbiased estimate of global infection frequency. Nonetheless, it is possible that other sampling biases, *e*.*g*., medical relevance of focal species, might impact our estimates if medically relevant groups have unusual infection frequencies relative to an “average” arthropod or nematode.

**Table 2.**
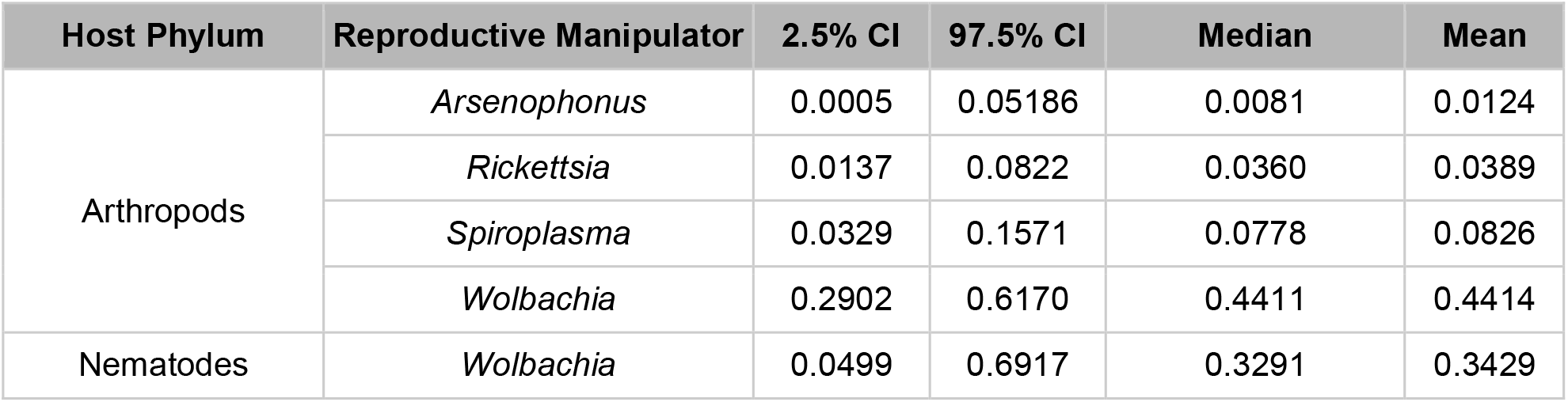
Estimated infection frequencies and confidence intervals from our data for *Wolbachia, Spiroplasma, Rickettsia* and *Arsenophonus* infecting arthropods and nematodes. All species in dataset were downsampled to maximum 100 individuals. We used a minimum threshold of 1 in 1000 of infected individuals within a species to classify a species as positively infected (Supplementary Method S3).

Among the other reproductive manipulators, we estimated *Arsenophonus, Rickettsia*, and *Spiroplasma* infected 1% (95% CI 0-5%), 3% (95% CI 1-8%), and 8% (95% CI 3-15%) of arthropod species, respectively (Table 2). We were not able to estimate global infection rates of *Cardinium* because of the extremely low rate of positive infection in our dataset. Owing to the fact that we do not have positive and negative controls readily available for each of these other reproductive manipulator clades, it is difficult to completely rule out infections that failed to map to the known reference(s) for each group and therefore induce a higher rate of false negatives. However, our results from mapping *Wolbachia* reads to extremely diverse reference genomes (e.g., 5-15% divergence) suggests that the rate of false negatives is low, provided the divergence within these other bacterial groups does not exceed our tested values and, as we note above, our raw frequency estimates are in line with previous work based on other methods [27, 48, 49].

### Taxonomic Distribution of Reproductive Manipulator Infections

The taxonomic distribution of *Wolbachia* spans 11 arthropod orders, out of the 47 tested (Figure 2). Across all of the arthropod species that we studied here, the orders with the greatest number of species sampled are Hymenoptera, Diptera, Lepidoptera, Coleoptera, and Hemiptera. These orders had 152, 195, 400, 139, and 91 species sampled, respectively. Positively infected samples are dispersed widely across the phylogeny, including hosts as distant as Coleoptera and Araneae. Horizontal transmission among species may contribute towards explaining *Wolbachia’s* large taxonomic distribution despite only modest sequence level divergence among *Wolbachia* strains.

**Figure 2.**
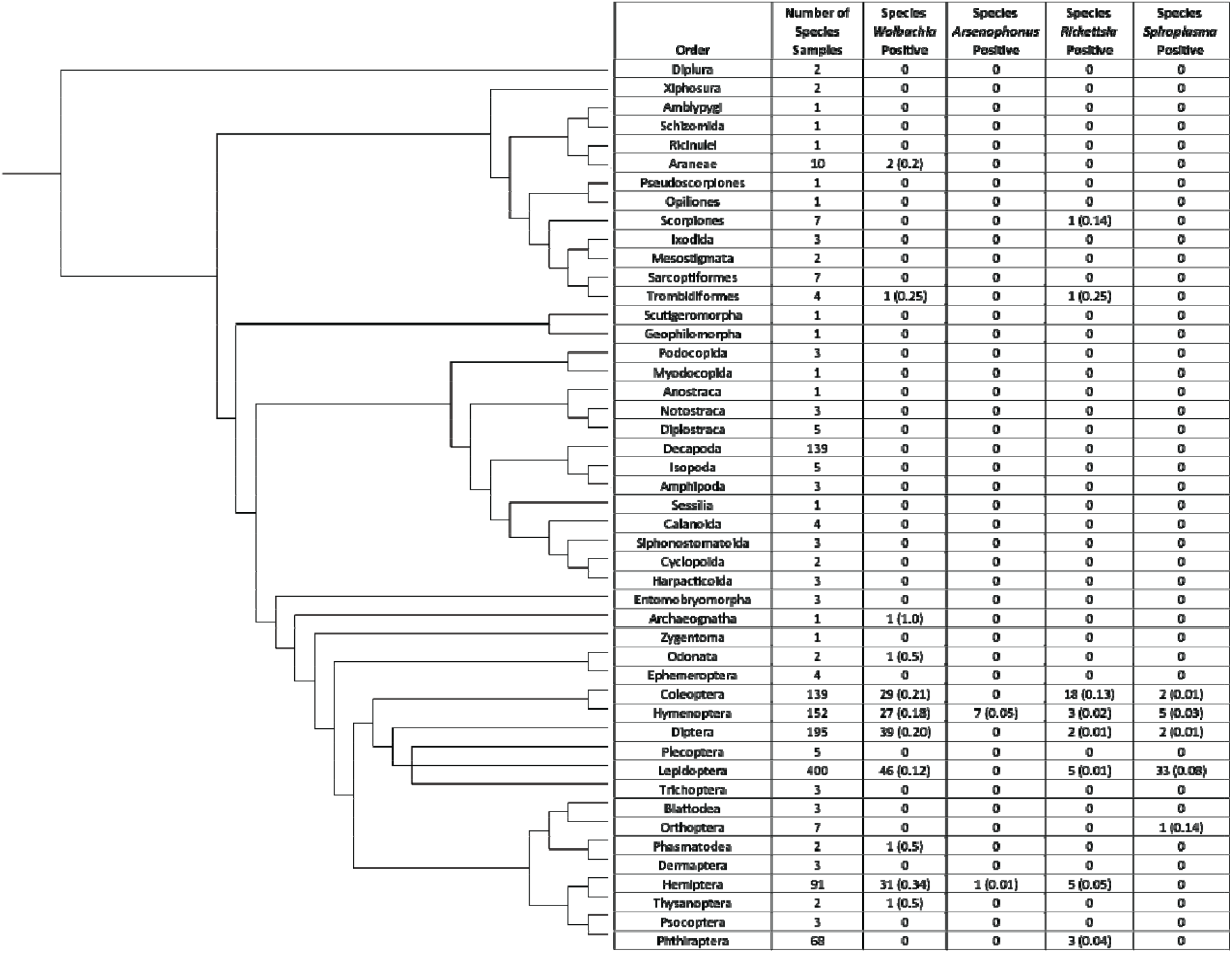
Phylogeny of Arthropoda orders tested and number of reproductive manipulator positive species within each order. The frequency of reproductive manipulator-positive species listed in parentheses. No frequency is listed if there was no infection within an arthropod order. We used the Tree of Life taxonomic and phylogenetic package and rotl <u>(Michonneau et al. 2016)</u>, to group host species by their orders.

*Wolbachia* infection frequencies vary substantially across insect orders. For example, Hemiptera, Hymenoptera, Diptera, and Coleoptera vary by approximately 30%, with 54%, 42%, 63%, 78% of species estimated to be infected, respectively (Table 3). Conversely, we estimated the lowest *Wolbachia* infection frequency for Lepidoptera (36%, 95% CI 9.28-70.4%, Table 3). Lepidoptera’s infection frequency, in particular, is slightly lower than the frequencies found in other orders of insects (*p* < 0.08, permutation test). Low frequencies might be the result of high fitness cost to infected hosts or from *Wolbachia’s* possible low transmission fidelity in these groups. Notably, *Wolbachia* infections feminize or kill males in Lepidopteran hosts [50]. Both of these phenotypes can impose fitness costs on populations harboring these symbionts because fewer males are available for mating and, in the case of feminization, males tend to prefer genetic females over feminized males, which have lower mating rates and receive less sperm [51]. Lower infection frequencies of bacteria with sex-ratio-distorting phenotypes might suggest feminization and male killing have a higher fitness cost, or that host populations can more easily subvert sex-ratio-distorting phenotypes compared to other reproductive phenotypes like cytoplasmic incompatibility and parthenogenesis. Sex-ratio-distorting phenotypes might therefore play a role in limiting infection polymorphism in host populations.

**Table 3.** *Wolbachia* global infection frequencies and confidence intervals generated for arthropod orders. All species in dataset were downsampled to maximum 100 individuals. Confidence intervals were generated using 1000 bootstrap replicates fitting a beta-binomial model to species infection frequency data among orders. A minimum infection frequency of 0.001 was used to classify a species as positively infected (Supplementary Method S3).

| Order | 2.50% | 97.50% | Median | Mean |
| --- | --- | --- | --- | --- |
| Coleoptera | 0.5272 | 0.9593 | 0.7940 | 0.7787 |
| Diptera | 0.3002 | 0.8983 | 0.6410 | 0.6333 |
| Hymenoptera | 0.1174 | 0.7727 | 0.4084 | 0.4236 |
| Hemiptera | 0.1719 | 0.9950 | 0.5118 | 0.5475 |
| Lepidoptera | 0.0928 | 0.7040 | 0.3503 | 0.3640 |

## Symbiont Titer

Replication control is important for vertically transmitted symbionts because the fitness of the symbiont is dependent on the fitness of the host. Reproductive manipulator infections must remain sufficiently high to ensure vertical transovarial transmission while being low enough to minimize pathogenic cost to a host [52]. Thus, replication control is important for reproductive manipulators to be maintained in host populations. However, the extent to which reproductive manipulator titers vary between orders of arthropods and among reproductive manipulator clades is largely unknown and might be an important component of the fitness costs reproductive manipulator species impose on their arthropod hosts.

### *In Vivo* and *In Silico* Validation to Estimate Symbiont Titer

To evaluate the accuracy of our titer estimation method (Figure 1F, 1G, 1E and Supplementary Method S4, Supplementary Table S13), we compared the relative density of *Wolbachia* within *D. melanogaster* and *D. simulans*, which are two of the most commonly studied species known to host *Wolbachia*. The titer of *Wolbachia* has been demonstrated to influence the degree of *Wolbachia’s* virulence on its host [53]. We hypothesized that titers of *Wolbachia* infecting *D. melanogaster* would be significantly lower than the titer of strains infecting *D. simulans* for three reasons. First, the strength of reproductive manipulation, which may be influenced by titer, is stronger in *D. simulans* than *D. melanogaster* [54, 55]. Second, the wRi strain commonly found in *D. simulans* has been reported to exhibit higher titers during embryogenesis than wMel does in *D. melanogaster* [56]. Third, wMel transinfected into *D. simulans* exhibited significantly higher titers than in its native host, *D. melanogaster [12]*.

Consistent with our expectations, we found that the strains inhabiting *D. simulans* exhibit significantly higher titer relative to those in *D. melanogaster* (*p <* 1e-05, one-tailed Mann Whitney U Test, Figure 3). In fact, for some extreme samples, titer in *D. simulans* shows an astounding >30:1 bacterial to host haploid genome complement ratio (Figure 3). Accounting for relative genome sizes, this indicates that *Wolbachia* can contribute approximately 30% as much DNA as the host does to the sequence data for that sample. To confirm these findings, we compared the amount of fluorescence due to *Wolbachia* DNA staining between stage 9/10a developing oocyte cysts from *D. melanogaster* flies infected with wMel and *D. simulans* flies infected with wRi, and found an average 2.1x more signal in *D. simulans* than *D. melanogaster* (*n*=15 and 19, respectively, Wilcoxon Rank-Sum *p* < 5.12e-05; Figure 3, Supplementary Method S5). Thus, the bioinformatically predicted titer differences between *Wolbachia* strains in *D. melanogaster* and *D. simulans* are also reflected using a completely orthogonal method to assay titer in the female germline. Importantly, this suggests that our *in silico* approach can yield information about *Wolbachia* abundance in tissues that are most relevant to understanding *Wolbachia* transmission (i.e., the female germline).

**Figure 3.**
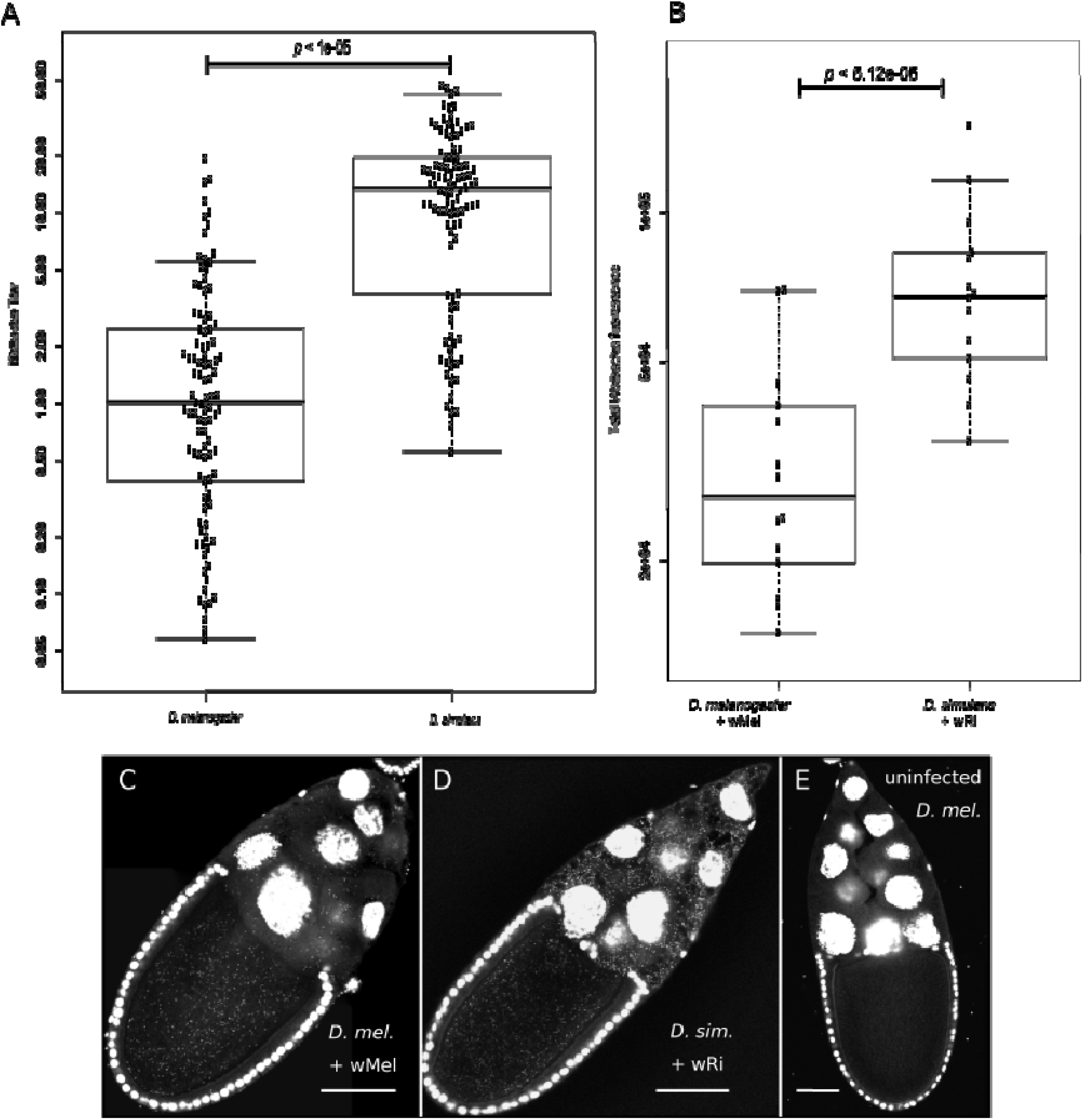
Wolbachia titer among *D. melanogaster* and *D. simulans* computed from our **(A)** in silico titer estimate approach (y-axis log10 scaled) and **(B-E)** in vivo fluorescence assay in stage 10a oocyte cysts. **(B)** Quantification of *Wolbachia* fluorescence, showing a significant difference in titer between host species (Wilcoxon rank sum test). **(C)** *D. melanogaster* and **(D)** *D. simulans* imaged propidium iodide DNA staining and confocal microscopy. **(E)** *Uninfected D. melanogaster* oocyte shows no evidence of *Wolbachia* staining. Scale bars = 50um.

### Comparative Study of Titer Across Symbiont Taxa

While symbiont titer does not necessarily predict the type or intensity of reproductive manipulation phenotypes in insect hosts [53], it does generally predict the virulence, or cost of infection, to a host [13, 14]. High titers are necessary to ensure an adequate number of bacteria make it to the germline for reliable vertical transmission. However, too high of titer may negatively impact host fitness. Since endosymbiotic bacteria like *Wolbachia* are maternally inherited, strong fitness costs to the host would also impact the fitness of *Wolbachia*. Therefore, evolutionary theory predicts symbiont titer will evolve towards a “goldilocks” zone, minimizing fitness costs to host and maximizing vertical transmission of symbiont bacteria [57]. The mechanism for this could either be via selection on symbiont growth or physiological activity [58, 59] or selection on host sanctions and regulatory mechanisms employed on the symbionts [14, 15, 59, 60].

We sought to identify the largest contributing factors to symbiont titer variation. Variation in titer may be a consequence of genetics (including host, symbiont, and combined host-symbiont genotype interactions) or a result of other biological variation among genetically similar individuals (*e*.*g*., due to ecology). We found significant inter-sample variability within species (Figure 4A, Supplementary Table S14). For example, there is a more than 200-fold range in *Wolbachia* titer across *Drosophila melanogaster* samples, consistent with previous estimates of titer in another *Drosophila* species *[61]*. Similarly, our results are in the range of previous estimates of *Wolbachia* titer in *Aedes Aegypti [62]*. To determine the degree to which host species influences variation in titer, we fit a generalized linear model to the data (Supplementary Table S14). We found that *Wolbachia* titer in arthropods varied significantly between arthropod species (*p <* 2e-16, GLM, Figure 4A), suggesting a strong genetic component shaping symbiont titer among arthropod species. Indeed, we found that this model can explain 36% of variation in titer. The remaining 64% of titer variation may be explained by within species differences, such as inter-individual variability and uncontrolled extrinsic factors, such as local diet or habitat. Our results therefore support the idea that symbiont titer varies considerably both within and between species.

**Figure 4.**
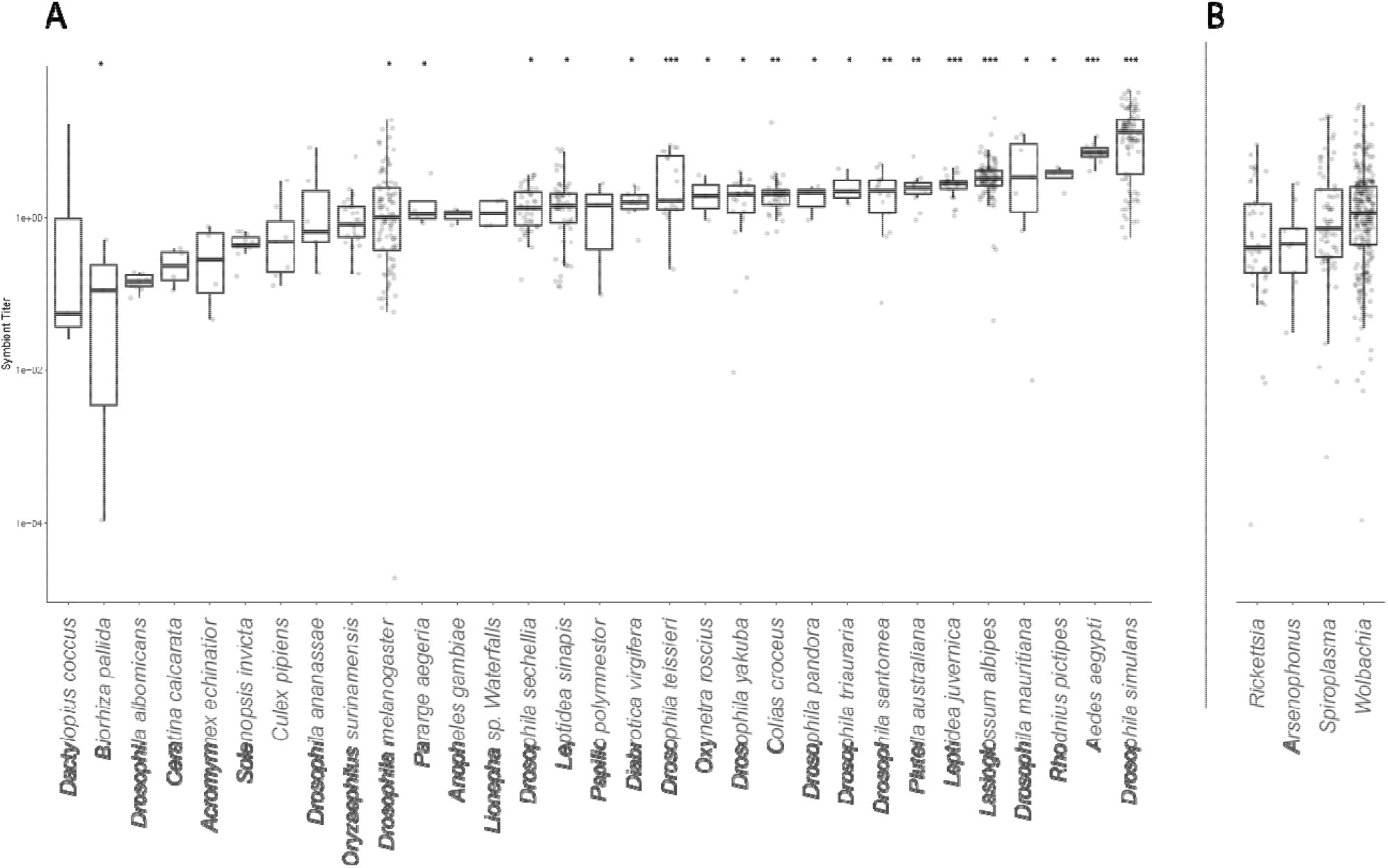
Titer variation across species infected with *Wolbachia* and between diverse reproductive manipulator clades. To increase readability of both plots, categories were randomly downsampled to show 100 samples. The y-axis is log10 scaled. **(A)** Titer for *Wolbachia* positive arthropod samples were grouped by host species and plotted. Host species with at least three samples were plotted. **(B)** Titer for *Wolbachia, Rickettsia, Arsenophonus, Spiroplasma* infected samples. We plotted up to three samples for every species infected with *Wolbachia* to show the range of *Wolbachia* across tested arthropod species. These plots show us that titer variation within host species is significant, and this variation is not due to pooled sequencing samples. Our results suggest a symbiont and host genetic contribution to shaping within-host infection densities.

We hypothesize that *Wolbachia* titer in arthropod hosts could be significantly different compared to other reproductive manipulator bacterial titers because *Wolbachia* is found at substantially higher frequencies across arthropod species and many studies have reported that *Wolbachia* generally elicits smaller fitness effects on its hosts than do *Rickettsia, Arsenophonus*, and *Spiroplasma* [63–68]. Although an expanded model including reproductive manipulator as an explanatory variable provides a better fit to the data (*p* = 0.004, likelihood ratio test), this model explains only an additional 0.5% of the variance in symbiont titer relative to a model where only host species is included as an explanatory variable (see Methods). No reproductive manipulator clade significantly contributed to titer variation (*p >* 0.05, likelihood ratio test). Our results therefore indicate that titer does not systematically differ between clades of reproductive manipulators, contrary to our expectations based on their infection frequencies and pronounced differences in their phenotypic effects. Finding that *Wolbachia’s* titer is relatively consistent with that of other reproductive manipulators suggests that co-evolved symbiont titer may not contribute to its widespread host range. These results suggest that other co-evolutionary mechanisms, perhaps encoded by the host genome or regulating bacterial physiology, contribute to *Wolbachia*’s widespread taxonomic host distribution and generally decreased virulence relative to other widespread reproductive manipulator taxa.

If several individuals, some infected and some not, were pooled to produce a single library, this could affect titer estimates. Most concerningly, the SRA metadata is not always complete. To confirm that pooling does not strongly affect our estimated titer and therefore bias our results, we performed a literature search and annotated each sample as pooled or not pooled when this information can be confidently determined from the source publication (Supplementary Table S15). Then, we discluded samples that we could not discernably categorize. We fit a GLM model of the same form as above but with pooled sequencing status as an additional factor, we found that this model does not fit significantly better than a GLM fitting titer and arthropod host species (*p* = 0.4, likelihood ratio test). This suggests our estimates from the whole SRA are robust, and that pooling samples does not significantly impact the variation we see in titer across arthropod hosts in our dataset. Our approach therefore appears robust to vagaries associated with aggregating diverse and distinct public datasets for computational analysis.

### Host Genotypes Associated with *Wolbachia* Titer

Considering the strong influence *Wolbachia* strains can have on their hosts’ reproductive outcomes, it is likely that the host genome experiences strong selective pressures to respond to or control the bacteria. Indeed, previous work [14] identified a gene in the wasp host, *Nasonia vitripennis*, that effectively controls *Wolbachia* titer and has been under recent positive selection, likely since the wVitA strain horizontally transferred into the species. To address this question using our public data-sourced method, we used the DGRP to perform a genome-wide association study (GWAS) of titer in *D. melanogaster* using the DGRP GWAS webtool [26].

Although the present analysis is exploratory, there are several noteworthy results that may suggest mechanisms of host control over *Wolbachia* titer. We identified 16 candidate single nucleotide polymorphisms (SNPs) in the *D. melanogaster* genome associated with *Wolbachia* titer (Supplementary Table S16). One of the most strongly associated SNPs is found in a gene associated with the endoplasmic reticulum (ER) membrane (*XBP1*) [69]. Consistent with this functional prediction, a recent study on a wMel-infected *D. melanogaster* cell line found that *Wolbachia* resides within ER-derived membrane near and within the ER itself [70]. Modifying this membrane might therefore enable the host to impact *Wolbachia* titer on the cellular level. Additionally, another strongly associated SNP is found in a gene associated with actin binding and microtubule transport (CG43901). Recent work has shown that *Wolbachia* uses host actin for localization in host tissues [71, 72] and modifications to actin-binding proteins can clear *Wolbachia* infections in host individuals [73]. Finally, a SNP in CG17048 is also strongly associated with *Wolbachia* titer in the DGRP. This gene’s role in protein ubiquitination is consistent with previous results from a genome-wide RNAi screen that found this process contributes disproportionately to modifications of *Wolbachia* titer in cell culture [74]. These are therefore appealing candidate genes for evaluating the potential of natural host variation to control symbiont infections.

Functional work will be necessary to validate our specific predictions, nonetheless these titer-associated genetic polymorphisms suggest that the host genome is capable of evolving to control *Wolbachia* infections. Although the present association work is focused on a well-characterized genetic mapping panel in *D. melanogaster*, our results illustrate more generally the potential impacts of high throughput data mining for identifying both reproductive manipulator infections and candidate host genetic factors that may be involved in controlling the infections within host tissues.

## Conclusion

Here, we presented a reference-based approach to detect bacterial infections in short read data and highlighted the ways it can be used to generate insights into host-symbiont interactions. Indeed, our work is the first to estimate the global distribution of multiple reproductive manipulators across all sequenced arthropod and nematode hosts using full-genome high throughput methods. Moreover, we show that publically-available short read data can be used to interrogate other biological attributes of host-symbiont associations, such as titer. We found that symbiont titer, measured as relative genome copy number, is highly variable among arthropod host species, and that titer can vary 200 fold between and within host species. Furthermore, our database catalogs some of the vast phenotypic and genetic diversity of reproductive manipulators, which could be used directly as biological control agents to control the spread of infectious disease or indirectly to inform on the natural infection state of host targets.

While our approach relies on datasets gathered for a wide array of purposes and therefore requires a level of approximation, we have shown that accurate and robust predictions can still be obtained using this method. Moreover, as publicly available sequence data continues to accumulate at exceptional rates, this framework will become increasingly powerful relative to gathering purpose-built datasets to assay symbiont infection statuses and frequencies. More generally, our method and related approaches could be used to detect other microbial symbionts, such as medically relevant pathogens, or even viruses, for which a reference genome sequence is available. Hence, future work will build on this framework of leveraging increasingly vast datasets to conduct direct and precise hypothesis testing of fundamental questions in host and symbiont ecology and evolution.

## Acknowledgements

We thank the Corbett-Detig Lab and Sullivan Lab members for helpful feedback on this work. During this research, PM was supported by NIH training grant (T32 HG008345). This work was supported in part by an Alfred P. Sloan Fellowship and by NIH/NIGMS R35GM128932 to RC-D.

## Contributions

Conceived and designed research: PM, RC-D, SLR. Performed the computational screen: PM. Analyzed sequence data and developed software: PM. Oocyte imaging and analysis: SLR. Novel infection literature analysis: KA. Wrote the manuscript: PM, SLR, RC-D. Edited the manuscript: RC-D, SLR, PM.

