## Supplementary Methods S1-S8 for "Deep data mining reveals variable abundance and distribution of microbial reproductive manipulators within and among diverse host species"

**Supplementary Method S1 - Determining Infection Status using a BLAST-based approach**

*Downloading reads from arthropod and nematode samples*

First, we used fastq-dump 2.9.0 from the SRA Toolkit to download all from each sequencing run associated with each sample. For example, if a sample was sequenced in three runs, we would use six million reads to classify the infection status of that sample. An example of our command line is shown below.

$ fastq-dump --fasta --split-files -I --stdout -X 1000000 ERR1882042.sra > ERR1882042.fasta

*Computing summary statistics*

Next, we aligned these reads to the set of reproductive manipulator genomes using blastn (version 2.7.1). We computed three summary statistics to help describe the similarity between sample reads and reproductive manipulator reference genomes. These statistics were: 1) the breadth of coverage, 2) the variance coefficient, and 3) the expected depth of coverage, and are described below. To calculate these statistics, we first divided each reference genome into non-overlapping 5kb bins, and we placed reads into bins based on their reference match start position. If a read aligned to multiple places in a reference genome, only the match with the lowest e-score was considered.

We computed all summary statistics for each reference independently. First, we computed the breadth of coverage as the proportion of 5kb bins with at least one read. Second, the variance coefficient was computed as the mean variance of the number of binned reads across 5kb bins, normalized by the total number of reads in all bins. Lastly, we computed the expected depth of coverage by using the total base pair length of significant BLAST hits generated from our pipeline, the total number of possible reads in a sequencing run, and the number of reads sampled using our pipeline.

*Criteria for positive infection*

We determined a sample to be a candidate for a positive infection if it mapped to >90% of a reproductive manipulator reference genome and had an estimated coverage >1x. Homologous fragments from non-reproductive manipulator bacteria could be detected and would appear to look like low breadth of coverage data. An estimated coverage of 1x or greater was required to prevent low-frequency contaminants from producing potential false positives. If a sample had between 0.1 and 0.9 breadth of coverage, the full dataset was downloaded and the workflow repeated to update predicted infection statuses. This two-step procedure allowed us to substantially decrease runtimes, because 97% of samples can be categorized using just 2 million read pairs, and to avoid false negatives associated with low sequence coverage positive infections.

**Supplementary Method S2 - Validation of bacterial detection pipeline**

*Wolbachia Infection Status of the DGRP and Subsampling Efficacy*

To estimate the sensitivity and specificity of our method, we compared our results to the survey conducted by (Richardson et al. 2012), which focused on *Wolbachia* infections in *D. melanogaster*, using the Drosophila Genetic Reference Panel (DGRP)*.* Subsampling two million reads from each sequencing run generated no false positive nor false negatives (Supplementary Table S7). When sampling all reads instead, we found similar low error rates (Supplementary Table S8). This indicates our approach may be conservative for calling infections when read numbers are limited. Both of the false negative samples generated from the all reads experiment were determined positive for *Wolbachia* from PCR analysis, but were determined negative using a previous WGS analysis (Richardson et al. 2012). This conflicting infection status could have been caused by PCR sample contamination and subsequent false amplification of a *Wolbachia-*specific fragment. Similarly, low-level contamination during library preparations could result in false positives using our bioinformatic approach. However, this would have to be a significant amount of contamination, as fewer PCR cycles are run with general Illumina primers in most Illumina library preparation protocols. Therefore, due to the small amount of uncertainty associated with this validation set our estimated false-negative and false-positive rates should be viewed as conservative upper-bounds. These data suggest subsampling reads may lead to a sizeable fraction of false negatives that can be ameliorated by sampling all reads for ambiguous samples. To efficiently mine the SRA database, we used a hybrid method where we first classified samples using two million reads. The samples that fell with an intermediate breadth of coverage between 0.1 and 0.9 were then reclassified using all available sequencing reads.

*Excluded DGRP samples*

We excluded 16 DGRP samples from our analysis where four were not available on the SRA database, eight did not have corresponding PCR and WGS *Wolbachia* infection statuses, and four (DGRP303, DGRP307, DGRP774, and DGRP109) had discrepancies in infection status when using Freeze 1.0 and Freeze 2.0 sequencing data (Mackay et al. 2012). These four samples were classified by our pipeline as negative for *Wolbachia* infection using Freeze 1.0 data. However, when we considered Freeze 2.0 sequencing data (Huang et al. 2014), our pipeline classified these samples as positive for *Wolbachia.* We note that our results from Freeze 1.0 are consistent with Richardson et al. (Richardson et al. 2012), who also used these data. The discrepancy between infection statuses generated by Freeze 1.0 and Freeze 2.0 could be due to contamination of Freeze 2.0 sequencing data or dramatic changes in infection frequency within a line between sequencing library preparation. Due to this possibility, we excluded these four samples from our dataset.

*Effect of Divergent Reference Genomes*

Given that there may be unsampled genomic diversity among reproductive manipulator strains in nature, and this may bias our detection method, we characterized our method’s sensitivity to detecting divergent *Wolbachia* strains by comparing mapping results for a sample among diverse reference genomes (Figure 2, and Supplementary Figure S5)*.* In addition to wMel*,* we used reference *Wolbachia* strains from host *Culex quinquefasciatus, Onchocerca ochengi, Brugia malayi, Cimex lectularius,* and *Pratylenchus penetrans* from *Wolbachia* Supergroup B, C, D, F, and L, respectively. When compared to wMel, these genomes exhibit 5-15% pairwise sequence divergence within alignable regions. Impressively, our method never produces false positives when determining infection status across all references using two million reads per sequencing run (Supplementary Table S7 and S8).

False negative rates are similarly modest for all divergent Wolbachia references, at a maximum of 0-6% when aligning all reads in all but the most divergent reference genome (Supplementary Table S7). Across all arthropod samples, we expect most of the symbionts within these samples to have the most similarity to *Wolbachia* from supergroups A and B (wMel and wPip) (Casiraghi et al. 2005; Werren and Windsor 2000; Werren, Zhang, and Guo 1995). So, finding that these strains can still be detected when using a nematode *Wolbachia* strain reference suggests that our method is very sensitive and the majority of *Wolbachia* infections were detected. Accuracy data for each reference genome are available in Supplementary Table S7 and S8. Taken together, these data indicate that subsampling sequencing read data is a computationally efficient method to determine reproductive manipulator infection statuses of the majority of host samples (*i.e.*, 97.5% can be accurately classified using a subsample of two million reads).

*Comparison to other methods*

A previous study by (Pascar and Chandler 2018) used a related bioinformatic pipeline to screen arthropod individuals for *Wolbachia* infection. The key difference between our method and theirs is that they selected only three loci (*wsp, ftsZ,* and *groE* operon sequences) rather than complete genomes, and required extremely similar blast hits to categorize samples as positive. (98 bp BLAST length matches at >= 98% identity to one or more of the reference genes and three or more matching sequence reads). They screened 2,545 arthropod sequencing runs, a subset of what we consider here, and found 173 (6.8%) of them were candidates for a positive *Wolbachia* infection which composed 11.8% of species tested. Their estimate ranges on the lower spectrum of *Wolbachia* prevalence estimates. Therefore, we ran our classifier on sequencing runs that Pascar’s method determined to be uninfected in order to compare the sensitivity of their method using our whole genome approach.

Of the 2365 runs classified as negative for *Wolbachia* infection, we found our method classified 22 sequencing runs as positive (Supplementary Table S5). Considering the median breadth of coverage and estimated coverage depth was 0.98 and 11x, respectively, we are confident that these sequencing runs are strong candidates for positive *Wolbachia* infection. If we are to consider these as positive samples, we estimate that the false negative rate of Pascar and Chandler’s method is 11%. Given that their approach is relatively conservative in identifying positive infections, it is unsurprising that we find a low proportion of false positives, which at 1.7%, is similar to our own.

*Identifying divergent Wolbachia strains*

To further confirm the accuracy of our approach across a broader phylogenetic sampling of *Wolbachia* strains, we estimated the infection status for known positive infections of *Zootermopsis nevadensis* (termite), *Ctenocephalides felis* (flea)*, Folsomia candida* (springtail)*,* and *Osmia caerulescens* (solitary bee) using our BLAST-based approach. All of the sequencing runs were previously determined to harbor *Wolbachia* live infections (Gerth et al. 2014) and classified into supergroup H, B, E, and F, respectively. Importantly, some of these divergent supergroups are not represented in our reference database and therefore represent the most challenging cases for accurate detection using our pipeline. All host samples were found to be positive for *Wolbachia* infection except for a sample from supergroup H, even after running our pipeline on all the reads from that sample (Supplementary Table S4). These results suggest that our method can detect *Wolbachia* from genetically divergent supergroups, but may not detect all of *Wolbachia’s* genomic diversity. Future efforts may mitigate this challenge by incorporating an increasingly diverse array of reproductive manipulator reference genomes.

We note that other reproductive manipulators may also pose other challenges for accurate quantification beyond the specifics that we encountered in *Wolbachia*. However, in the absence of a high quality validation set and numerous true positives which are available for *Wolbachia*, it is challenging to formally evaluate our pipeline’s performance in the other reproductive manipulators. Nonetheless, the overall robustness and accuracy when applied to *Wolbachia* positive and negative controls, and the consistency of our frequency and titer estimates with previous results suggests that there are few significant biases associated with our approach.

**Supplementary Method S3 - Beta-binomial model fitting**

*Beta-binomial rationale*

Our study aims to directly sample and determine the infection statuses of more animals of any study to date. However, there is an inherent sampling bias in any study that tries to estimate the prevalence of reproductive manipulators within species where infection might occur at intermediate or low frequencies. For example, the probability of sampling one individual from a population with high infection frequency (i.e. many individuals are infected) is higher than sampling an infected individual from a population with low infection frequency. The probability, then, of classifying a population as infected is dependent on the number of individuals tested and the frequencies of infection within each species. To evaluate and correct for this ascertainment bias, we use a beta-binomial distribution to estimate the total proportion of reproductive manipulator infected species.

*Mitigating sampling bias by downsampling*

We tested whether pruning our dataset was necessary to estimate the global infection frequency of reproductive manipulators in arthropods and nematodes. Since the beta-binomial model is positively influenced by the number of species, and number of individuals sampled per species in the dataset, Hilgenboecker *et al.,* 2008 tested a variety of downsampling methods in order to mitigate the influence of a non-uniform sample set would have on their global estimates. In addition, Hilgenboecker *et al.,* 2008 gathered their infection status data from studies measuring the prevalence of *Wolbachia.* The authors state that curating a sample set from these studies almost certainly introduced bias towards finding *Wolbachia* in a sample, as the studies focused on species known, or at least suspected, to have *Wolbachia.* Our method contrasts with this sampling approach, and implements a more random strategy to sampling arthropod and nematode species. Although there is still a sampling bias to what species are sequenced (i.e. medically relevant, model systems), our approach does not specifically bias toward species that are thought to harbor reproductive manipulators (Supplementary Table S8).

To test the effect of non-uniform sample size distribution among host species, we fit beta-binomial models to downsampled datasets. To downsample, we chose a maximum threshold (nj_max) in which all species would be downsampled to have a maximum of nj_max individuals. To downsample individuals within a species to nj_max, we randomly chose individuals without replacement from within the species. Confidence intervals were computed via 1000 bootstraps replicates.

With the full dataset, the *Wolbachia* global infection frequency 95% confidence interval is estimated to be between 0.3 and 1 (Supplementary Figure S2). Estimates of *Wolbachia* global infection frequency decrease as datasets were downsampled. Moreover, we see the 95% confidence interval become tighter around the mean as we downsample the original dataset (Supplementary Figure S2). These results taken together suggest that the beta-binomial model is influenced by the few large sample sizes in the dataset and varying global infection frequencies can be produced depending on the data set used. Nonetheless, we see a stable global infection frequency of *Wolbachia* between 0.4 and 0.5 when species are downsampled to 100 individuals. Therefore, we computed global infection frequencies using a dataset downsampled to 100 individuals per species.

**Supplementary Method S4 - Estimating symbiont titer**

We used DNA reads to estimate the ratio of symbiont genome compliments to host genome compliments, hereafter referred to as titer. We estimated the number of symbiont genomes by using our method described above with an expanded blast database including all symbiont genome assemblies on NCBI (Supplemental Table S1). In total, the symbiont database used to estimate titer had 141 symbiont genome assemblies. We used a set of single copy orthologous proteins from arthropods using OrthoDBv9 (Zdobnov et al. 2017) to estimate the density of host cells in a sample. This set contained 1066 proteins, from 133 taxonomic groups spanning Arthropoda, which amounted to a total 312,654 amino acids of reference sequence (Zdobnov et al. 2017). We used a tblastn based approach to locally align host nucleotide reads to the arthropod orthologous protein sequences. Because not all arthropod single copy ortholog proteins might be present in the host sample, or because there might be regions of the protein that are divergent from the host sample, we used the average of maximum depth across orthologous proteins that have reads mapping to them as an estimate of host titer. We corrected for the ploidy difference between host (assumed to be diploid) and symbionts (haploid) by halving the computed host coverage to result in a titer estimation (symbiont haploid : host pseudo haploid).

We confirmed that our tblastn approach to estimate host genome coverage is consistent with a reference-based alignment approach using a set of single copy orthologous genes representing arthropod genetic diversity (Spearman’s rho = 0.81, Supplementary Figure S3). We also compared our tblastn approach to a reference based approach using whole genome sequencing to estimate titer in DGRP samples (Spearman’s rho = 0.95, Supplementary Figure S4) (Richardson et al. 2012). These results corresponded well to the results from our ortholog approach, so we continued with our previously described workflow to estimate host coverage.

Furthermore, our titer estimates might be impacted by the distance to the reference selected. We tested this potential impact by selecting the reference with the second highest read coverage using BLAST. We would expect host coverage estimates to vary significantly between references if our titer estimates were impacted by the distance to the reference selected. For each positively infected sample in Figure 4, we compared its coverage to the closet reference and second closest reference. We did this for all 132 *Spiroplasma, Rickettsia,* and *Arsenophonus* samples in Figure 4 and found no major differences (Supplementary Table S13, Supplementary Figure S11). This suggests that our method is robust in estimating stable symbiont genome coverages especially from Wolbachia Supergroups A and B, of which comprise the majority of tested Arthropod infections (Supplementary Table S13, Supplementary Figure S11).

**Supplementary Method S5 - Drosophila oocyte sampling, imaging and analysis**

*Markers and balancers*

The *D. melanogaster* stocks used were carrying the markers and balancers w[1]; Sp/Cyo, Sb/Tm6, Hu or the germline double driver: P{GAL4-Nos.NGT}40; P{GAL4::VP16-Nos.UTR}MVD1. These stocks were infected with the wMel strain of *Wolbachia.* The *D. simulans* stock used was w[-] and was infected with the wRi strain endogenous to *D. simulans* populations in North America.

*Collection*

Flies were collected shortly after eclosion and transferred to new white food for three to five days. We dissected the ovaries of approximately 10 flies of each species in 1xPBS, and “fluffed” the ovarioles with pins to separate them. The ovaries were fixed in formaldehyde and heptane, RNAse A treated, and stained with propidium iodide, as described in (Russell et al. 2018).

*Imaging and analysis*

We mounted the stained oocytes on glass slides and imaged them with a SP5 Leica confocal microscope using a 63x objective. We imaged oocytes through their middle planes, taking optical sections every 0.38 um, the Nyquist value. For analysis, we picked comparable planes approximately halfway through the oocyte for all imaged oocytes, and created 3D brightest point projections from three slices, representing the diameter of one *Wolbachia* cell (~1 𝜇m), in ImageJ. We analyzed these images for fluorescence due to *Wolbachia* by manually removing fluorescence due to host cell nuclei, thresholding the image to eliminate background noise, and measuring the fluorescence contained within the entire oocyte cyst as described in Russell et al. 2018. We calculated the corrected total cell fluorescence (CTCF) for each cyst with the following formula: CTCF = Integrated Density–Area of selected cell X Mean fluorescence of background readings. Results were plotted with the vioplot package in R.

**Supplementary Method S6 - Identifying previously unreported infections**

We searched the literature to identify previously unreported *Wolbachia, Spiroplasma, Rickettsia,* and *Arsenophonus* infections in our SRA scan dataset. We used Google Scholar, and searched [species] + [reproductive manipulator name]*.* If no published results were found, we also searched the results of (Pascar and Chandler 2018) who did an SRA scan, albeit smaller than ours. If no infection was found using both these methods, a species was determined to be a novel infection. Our method to determine novel infections is not exhaustive, especially if we consider species nomenclature can change over time. Nonetheless, these candidate novel infections illustrate the potential impacts of our method.

**Supplementary Method S7 - Co-infection permutation test**

To test whether our observations of co-infected species, where one species harbors observations of two or more reproductive manipulator strains, exceeded what would be expected by chance, we conducted a permutation test. We also performed a similar test for individual-level coinfections, where we asked if the number of individuals that were coinfected in our data exceeded what would be expected by chance. To do these, for each species and for each reproductive manipulator we drew a probability, *p,* from the beta-binomial distribution of each reproductive manipulator infection frequency. Then, for each species, we drew samples from a binomial distribution with parameters *p* and *n* where *n* is downsampled to a maximum of 100 for each species for consistency with our analyses*.* We counted the number of individuals with co-infections as well as the number of species with co-infection and reported a *p-*value based on how many of the 1000 bootstrap replicates were greater than or equal to our observed counts (Supplementary Figures S9 and S10). This approach is preferable to an individual-based permutation because it controls for the autocorrelation among individuals within a population by requiring that the infection frequency be fixed for each population or species. It is therefore unaffected by differences in sample sizes among species.

**Supplementary Method S8 – Titer literature review and controlling sampling biases**

Variation in tissues and pre-sequencing treatment of samples could affect symbiont titer levels. Additionally, pooling several individuals for DNA extractions might also impact mean titer estimates if infections are polymorphic within species. Considering this bias, we reviewed and aggregated the methods used to sequence samples in our comparative study of titer among symbiont species (Figure 4, Supplementary Table S15). We used the project accession numbers found in the SRA metadata to trace back to primary literature describing the origin of each sequencing dataset. We report and retain samples that we could confidently categorize as pooled on not-pooled. We fit a generalized linear model to test for effects of pooling on titer variation in our arthropod dataset (See “Comparative Study of Titer Across Symbiont Taxa” in Results, Supplementary Table S15).
