## Supplementary Figures S1-S11 for "Deep data mining reveals variable abundance and distribution of microbial reproductive manipulators within and among diverse host species"

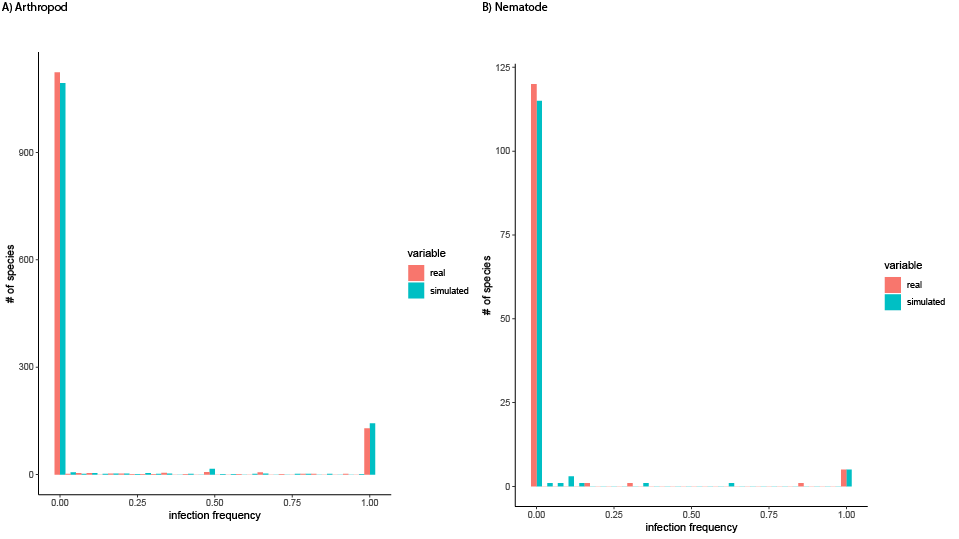

**Figure S1**. Analytical data from the SRA scan (real) and simulated infection frequencies (simulated) for **(A)** arthropods and **(B)** nematodes showing a beta-binomial distribution. Dark bars show empirical data and light bars show simulated infection frequency data from the beta-binomial model fit to our observed frequency data. The beta-binomial model was fit to *Wolbachia* infection frequencies in arthropod and nematode species downsampled to 100 individuals. Beta-binomial parameters, prob and theta, were estimated to be 0.07 and 0.51 for arthropods, and 0.05 and 0.64 for nematodes.

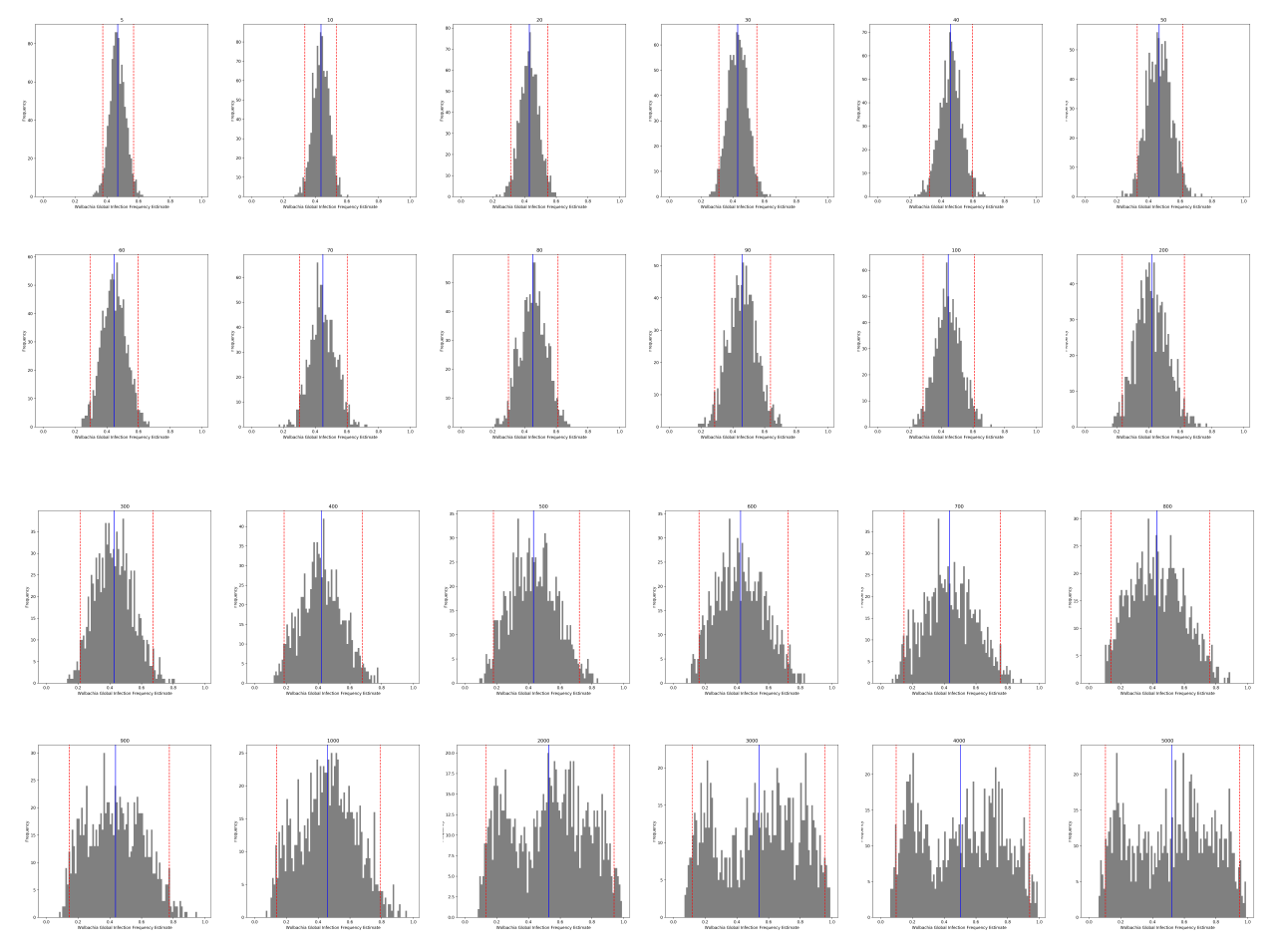

**Figure S2.** Bootstrap estimates of *Wolbachia* global infection frequency in insect species. We performed 1000 bootstrap replicates by sampling N insect species with replacement. Panel titles indicate the maximum number of individuals within an insect species (nj_max). If the number of individuals within a species (nj) exceeded nj_max, then we randomly sampled individuals with replacement until nj = nj_max. A beta-binomial model was fit to the downsampled insect data to estimate the global infection frequency of *Wolbachia* in insects. We used a minimum infection frequency threshold of 0.001 to classify a species as positive for infection. The mean frequency across all bootstrap replicates is indicated by the vertical blue lines, with 95% confidence intervals marked with dashed red lines.

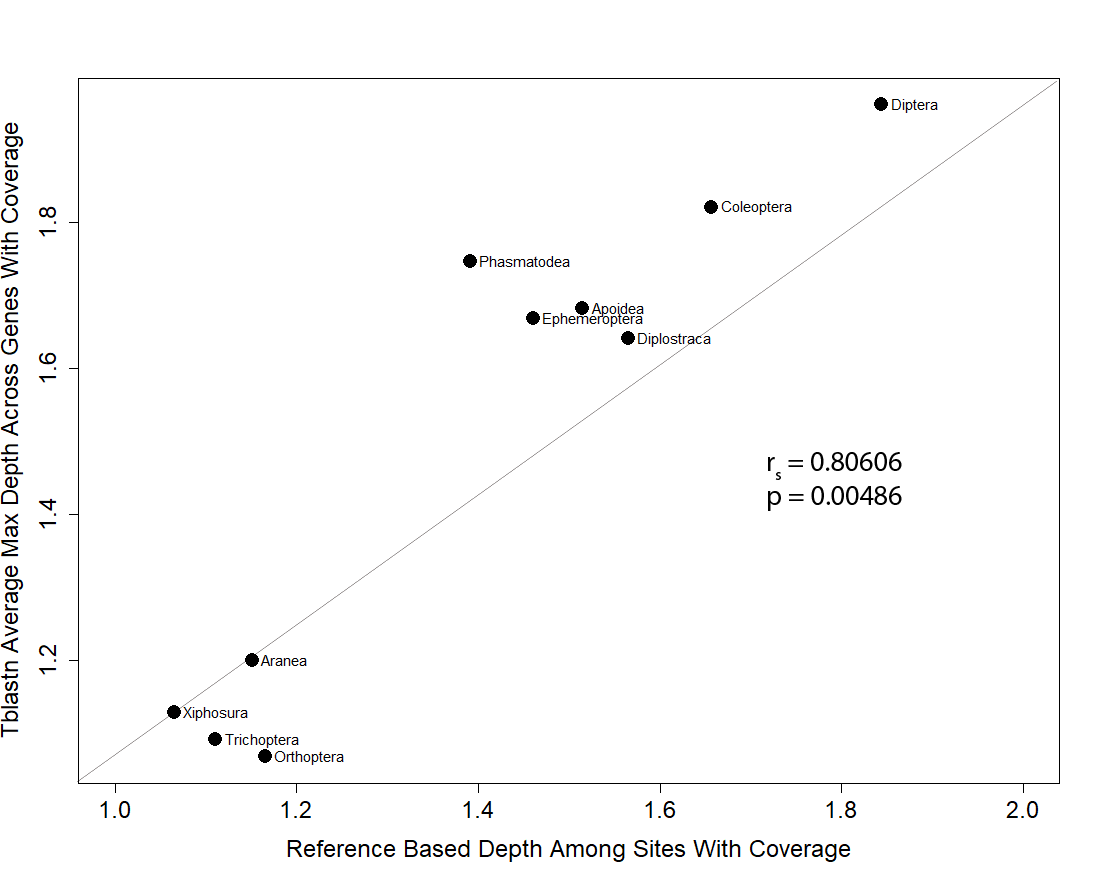

**Figure S3.** Comparison of titer estimation using single copy ortholog proteins (tblastn) versus a reference based approach (bwa mem). Estimation of arthropod host titer using ancestral orthologous protein sequences (y-axis) compared to depth of alignment to the reference genome (x-axis) (See Supplementary Method S7). Endosymbiont:Host titer was estimated using two million reads. Host titer was divided by two to generate a haploid titer comparison between *Wolbachia* and host (titer = *Wolbachia* 1C: host 1C). Correlation between our non-reference based approach to the reference approach suggest we can estimate titer across diverse host clades.

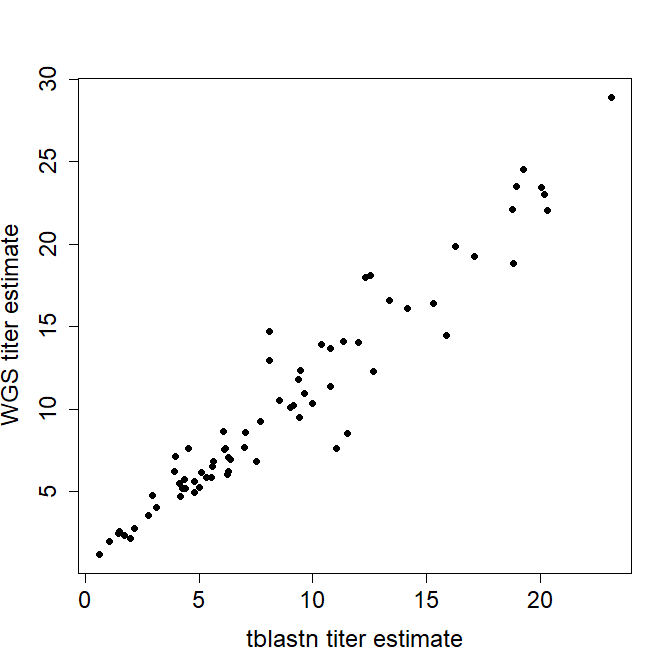

**Figure S4.** Estimation of W*olbachia* titer in *Drosophila melanogaster* from the DGRP using ancestral orthologous protein sequences (y-axis) compared to depth of alignment to the reference genome (x-axis) (See Supplementary Method S7). Two million reads were used estimate *Wolbachia:*host titer. Host titer was divided by two to generate a haploid titer comparison between *Wolbachia* and host (titer = *Wolbachia* 1C: host 1C). WGS titer was computed from the Richardson et al. 2012 data, and used as positive control. Spearman’s rho = 0.95, 2-tailed p-value = 9e-38.

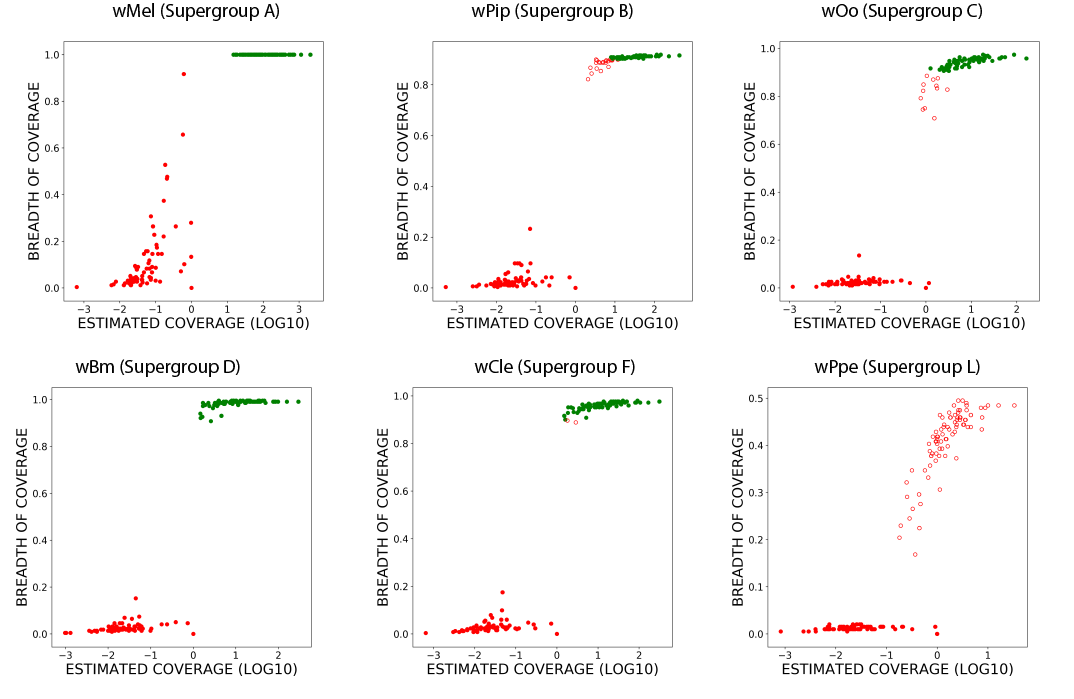

**Figure S5.** Using subsampled reads, our BLAST-based approach accurately determines the presence of *Wolbachia* for individuals from the Drosophila Genetics Research Panel when using all but the most divergent *Wolbachia* reference genomes. Two million reads were randomly subsampled from each sequencing run to produce these data. Each point represents a sample that was positively infected (green) or negatively infected (red) when mapped to the reference listed at the top of each plot. Circles are filled if the infection status predicted by our method matched the prediction determined by whole genome sequencing and PCR (Richardson et al. 2012). References listed from top left to right are *Wolbachia* from *Drosophila melanogaster* (supergroup A)*, Culex quinquefasciatus* (supergroup B)*, Onchocerca ochengi* (supergroup C)*, Cimex lectularius* (supergroup F)*, Brugia malayi* (supergroup D)*,* and *Pratylenchus penetrans* (supergroup L)*.* A sample was determined to have a *Wolbachia* infection if more than 90% of the 5 kb bins had at least one read and the estimated mean depth of coverage was at least 1x.

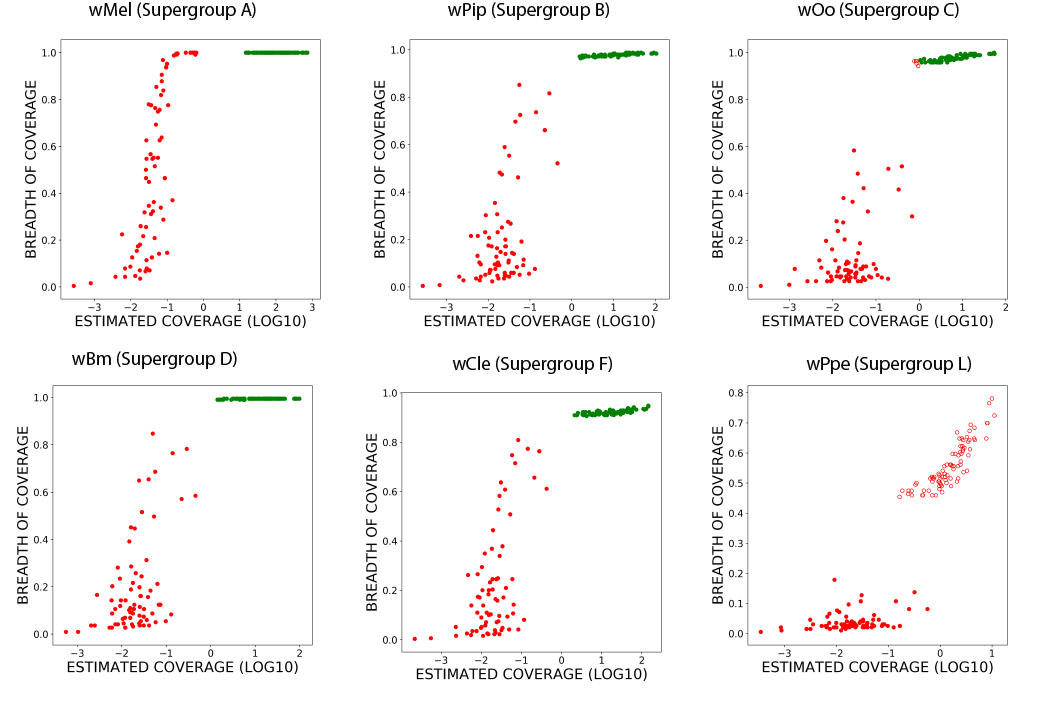

**Figure S6.** Using all reads, our BLAST-based approach accurately determines the presence of *Wolbachia* for individuals from the Drosophila Genetics Research Panel when using all but the most divergent *Wolbachia* reference genomes. Each point represents a sample who is positively infected (green) or negatively infected (red). Circles are filled if the infection status predicted by our method matched the prediction determined by whole genome sequencing and PCR (Richardson et al. 2012). References listed from top left to right are *Wolbachia* from *Drosophila melanogaster* (supergroup A)*, Culex quinquefasciatus* (supergroup B)*, Onchocerca ochengi* (supergroup C)*, Cimex lectularius* (supergroup F)*, Brugia malayi* (supergroup D)*,* and *Pratylenchus penetrans* (supergroup L)*.* A sample was determined to have a *Wolbachia* infection if there was more than 90% of 5 kb bins had at least one read, and an estimated mean depth coverage was at least 1x.

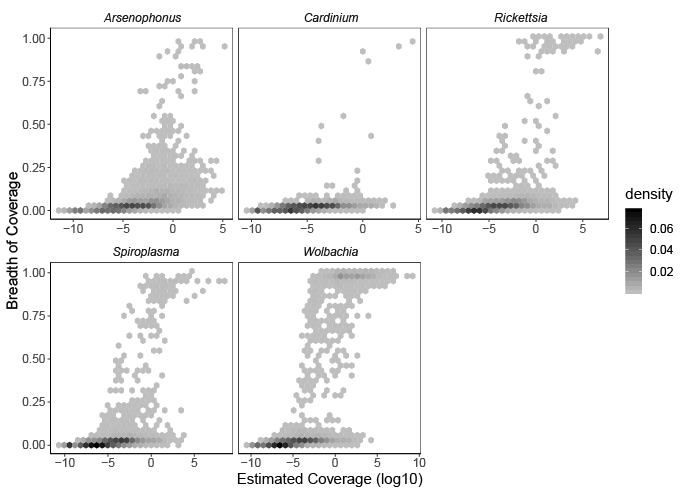

**Figure S7.** Summary statistics generated from the SRA arthropod scan. Each kernel density point is composed of arthropod sample DNA sequencing reads locally aligned to a reproductive manipulator reference genome. Reproductive manipulators are separated into panels. Samples with greater than 0.90 breadth of coverage and greater than 1x estimated coverage were considered positive (See Supplementary Method S1).

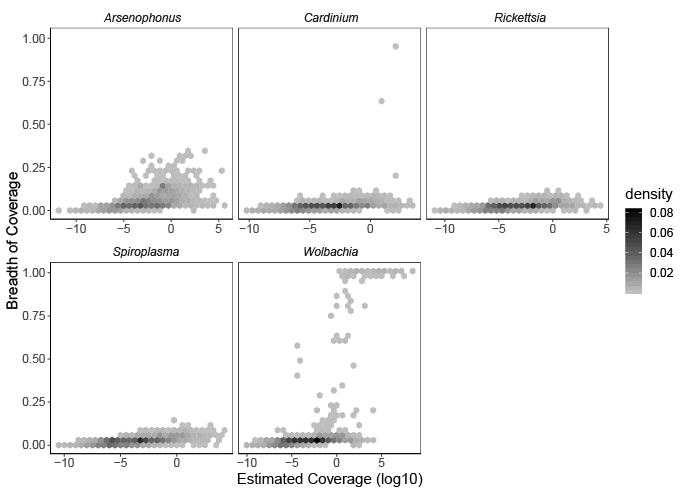

**Figure S8.** Summary statistics generated from the SRA nematode scan. Each kernel density point is composed of nematode sample DNA sequencing reads locally aligned to a reproductive manipulator reference genome. Reproductive manipulators are separated into panels. Samples with greater than 0.90 breadth of coverage and greater than 1x estimated coverage were considered positive (See Supplementary Method S1).

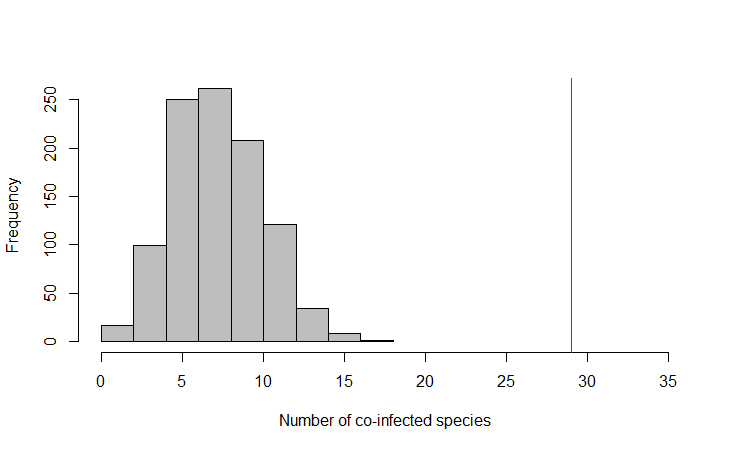

**Figure S9.** Permutation test results in fewer within-species co-infections than observed in empirical data (Supplementary Method S7). The red line indicates the number of co-infected arthropod species observed from the SRA scan.

.

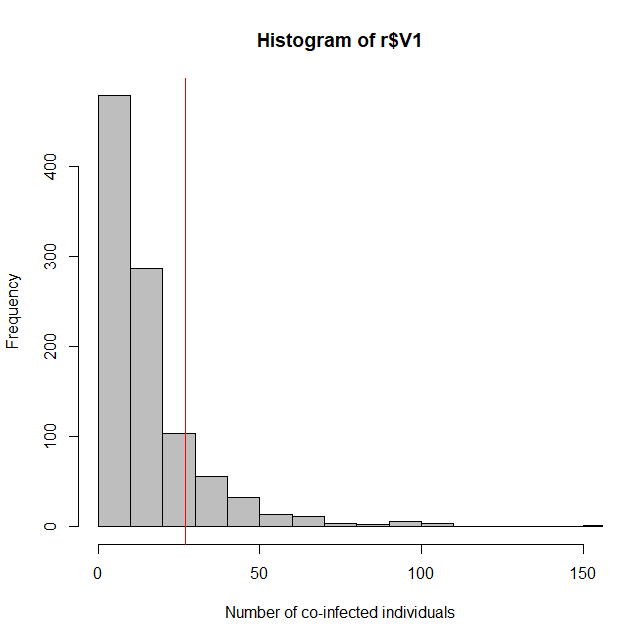

**Figure S10.** Permutation test results in fewer within-individual co-infections than observed in empirical data (Supplementary Method S7). The red line indicates the number of co-infected arthropod samples observed from the SRA scan.

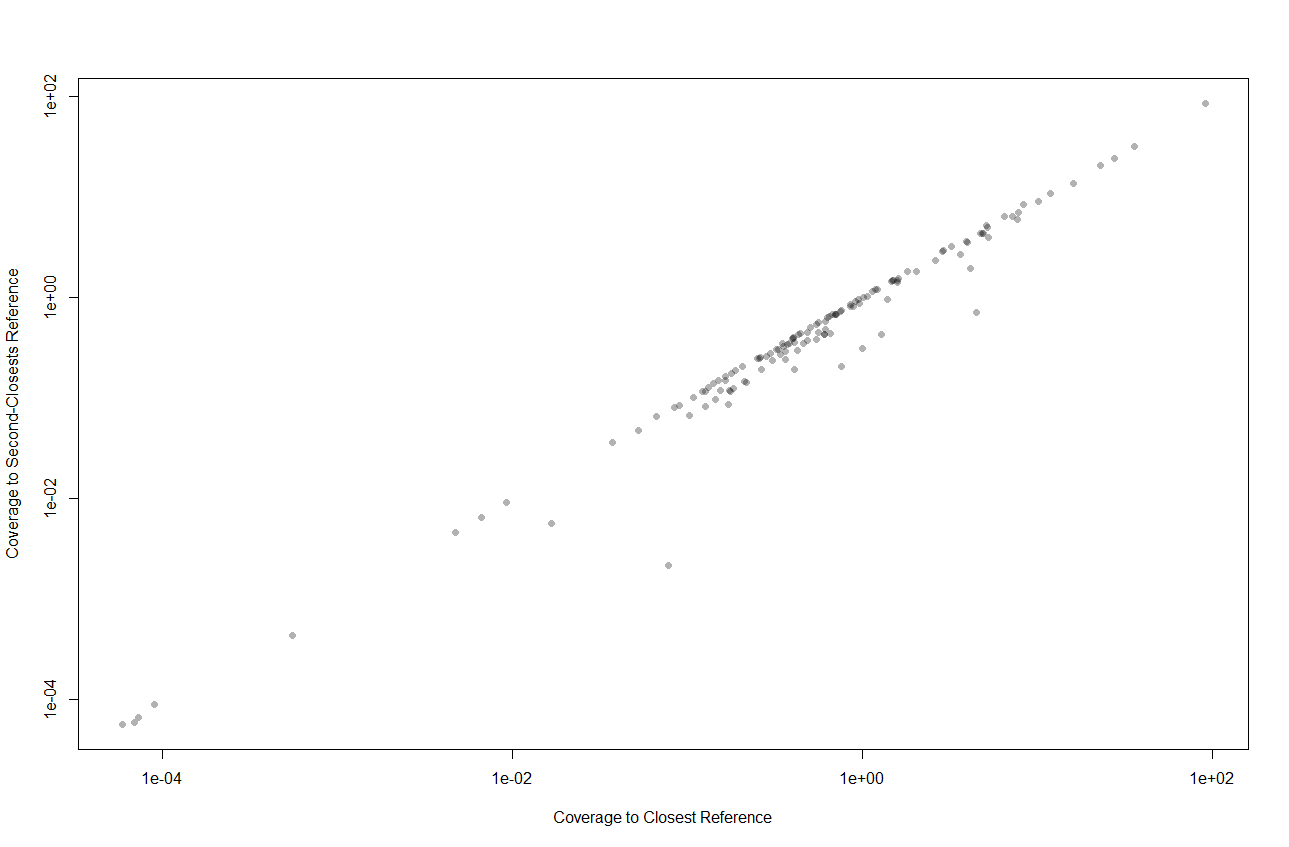

**Figure S11.** Comparison between the coverage (read depth) of the highest and second highest reference for each arthropod sample with a positive *Arsenophonus, Spiroplasma,* or *Rickettsia* infection. Our method estimates very similar symbiont coverages when using the best or second best references (Spearman’s rho = 0.98, *p* < 2.2e-16). These results suggest our method is robust in capturing consistent symbiont coverages. See Supplementary Method S4.
